# Evolutionary blocks to anthocyanin accumulation and the loss of an anthocyanin carrier protein in betalain-pigmented Caryophyllales

**DOI:** 10.1101/2022.10.19.512958

**Authors:** Boas Pucker, Nathanael Walker-Hale, Won C. Yim, John Cushman, Alexandra Crum, Ya Yang, Samuel Brockington

## Abstract

◻ The order Caryophyllales exhibits complex pigment evolution, with mutual exclusion of anthocyanin and betalain pigments. Given recent evidence for multiple shifts to betalain pigmentation, we re-evaluated potential mechanisms underpinning the exclusion of anthocyanins from betalain-pigmented lineages.
◻ We examined the evolution of the flavonoid pathway using transcriptomic and genomic datasets covering 309 species in 31 families. Orthologs and paralogs of known flavonoid synthesis genes were identified by sequence similarity, with gene duplication and gene loss inferred by phylogenetic and syntenic analysis. Relative transcript abundances were assessed to reveal broad-scale gene expression changes between betalain- and anthocyanin-pigmented lineages.
◻ Most flavonoid genes are retained and transcribed in betalain-pigmented lineages, and many also show evidence of extensive gene duplication within betalain-pigmented lineages. However, expression of several flavonoid genes is reduced in betalain-pigmented lineages, especially the late-stage genes dihydroflavonol 4-reductase (*DFR*) and anthocyanidin synthase (*ANS*). Notably flavonoid 3′,5′-hydroxylase (*F3′5′H*) homologs have been repeatedly lost in belatain-pigmented lineages, and Anthocyanin9 (*AN9*) homologs are undetectable in any betalain-pigmented lineages.
◻ Down-regulation of *ANS* and *DFR* homolog expression (limiting synthesis) and reiterative loss of *AN9* homologs (limiting transport), coincident with multiple shifts to betalain pigmentation, are likely crucial the loss of anthocyanins in betalain-pigmented Caryophyllales.

## INTRODUCTION

Plants produce a vast array of specialized pigments generating different colours (Li *et al*., 1993; Last, 2019). Pigments are involved in a huge number of critical biological functions including photosynthesis, pollination, fruit and seed dispersal, and the protection against abiotic and biotic stress (Demmig-Adams *et al*., 1996; Tanaka *et al*., 2008). Plant pigments are classified into several major classes based on their biochemical structure and synthesis: chlorophylls, carotenoids (carotenes, xanthophylls), flavonoids (anthocyanins, proanthocyanidins, flavones, flavonols) and betalains (betaxanthins, betacyanins) (Winkel-Shirley, 2001; Tanaka *et al*., 2008; Timoneda *et al*., 2019). Many of these pigment classes such as chlorophyll, carotenoids and flavonoids are essentially ubiquitous across land plants, but notably, some pigment classes have occasionally been lost in selected lineages, for example, the loss of chlorophyll in holo-parasitic lineages, and the repeated losses of flavonoid-derived anthocyanins in multiple lineages within the flowering plant order Caryophyllales (Bate-Smith, 1962; Mabry & Turner, 1964; Molina *et al*., 2014).

In Caryophyllales, an unusual class of pigments, the betalains, replace the otherwise ubiquitous anthocyanins. In the betalain- pigmented species of Caryophyllales, anthocyanin pigmentation has never been detected (Bate-Smith & Lerner, 1954; Mabry & Turner, 1964) and, conversely, the anthocyanic lineages within Caryophyllales do not produce betalains (Clement & Mabry, 1996). Based on these data, it has been proposed that anthocyanins and betalains are mutually exclusive (Stafford, 1994; Clement & Mabry, 1996). However, betalain-pigmented Caryophyllales continue to maintain flavonoids like flavonols, and proanthocyanidins in the seed coat (Shimada et al., 2005). The phylogenetic distribution of anthocyanin and betalain-pigmented lineages is homoplastic, with multiple betalain-pigmented clades sister to anthocyanin lineages (Sheehan *et al*., 2020) (**Fig. 1**). This interdigitated pattern of betalain and anthocyanin-pigmentation has traditionally been explained by an origin of betalains early in Caryophyllales followed by multiple reversals to regain anthocyanin pigmentation **(Fig. 1a)**. However, more recent evidence suggests that the betalain synthesis pathway arose multiple times within Caryophyllales (Sheehan *et al*., 2020), which in turn implies multiple independent losses of anthocyanins (**Fig. 1b**). In a scenario of multiple shifts to betalain pigmentation, loss of anthocyanin pigmentation is implied to be less readily reversible, with less scope to invoke subsequent reversals back to anthocyanins from a betalain-pigmented ancestor, in contrast to traditional explanations (**Fig. 1b**) (Brockington *et al*., 2011, 2015).

**Figure 1.**
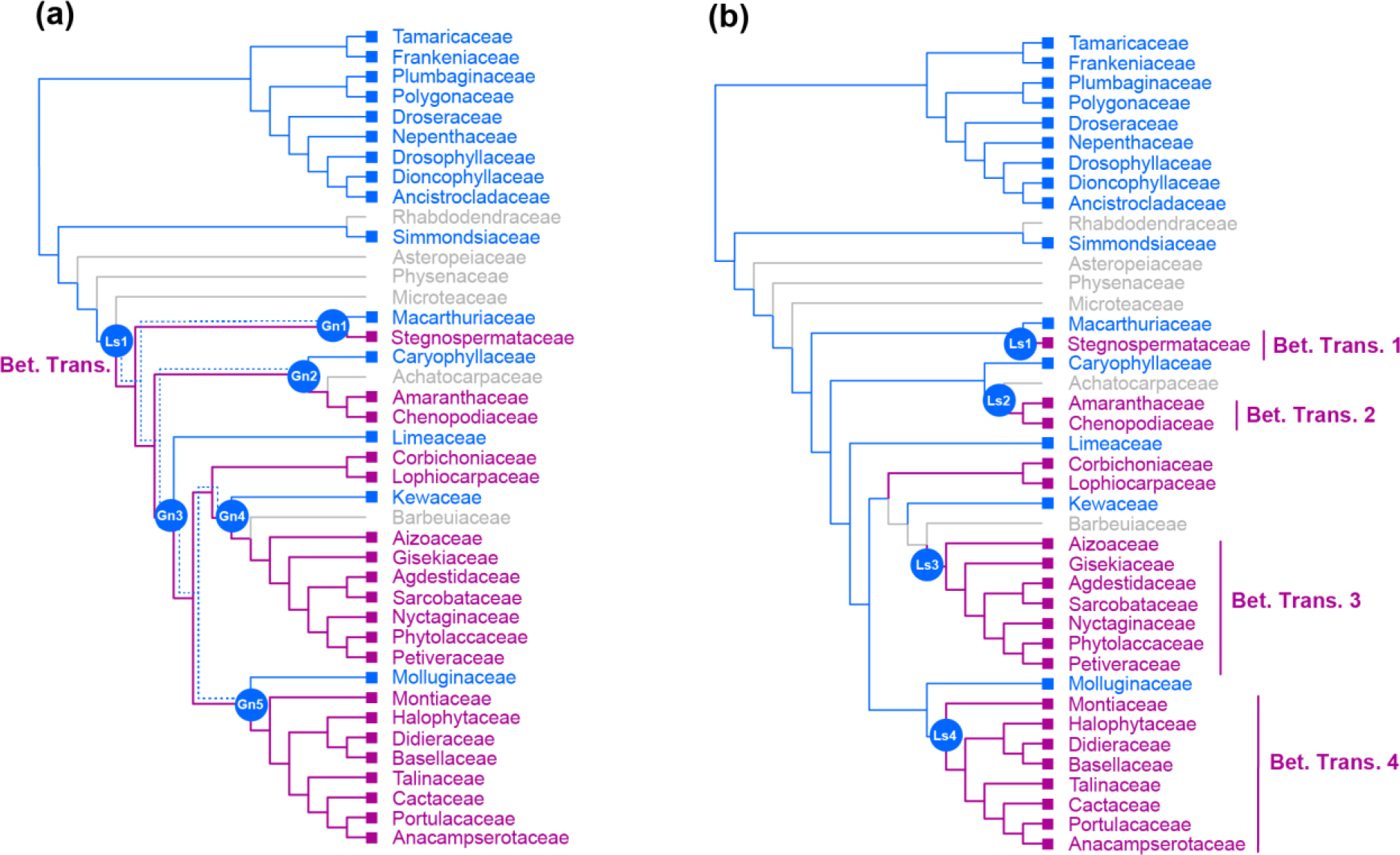
Two alternative hypotheses of pigment evolution in Caryophyllales. (a) a single origin of betalain pigmentation (*sensu* Brockington *et al*., 2015) implies a single loss of anthocyanins and subsequently five independent reversals (Gn1-5) back to anthocyanin pigmentation (dotted blue lines represent maintenance of anthocyanin pathway genes); (b) in this scenario all instances of anthocyanin pigmentation represent retention of the plesiomorphic state, and multiple transitions (Bet. Trans 1-4) to betalain pigmentation (se*nsu* Sheehan et al., 2020) implying at least four independent losses of anthocyanin (Ls1 -4). Blue=anthocyanin, pink=betalain, grey=unknown. Tree topology and color coding based on the mutual exclusion between the betalain and anthocyanin pigmentation and the family level phylogeny of Sheehan et al., 2020.

Anthocyanin pigmentation requires biosynthesis of the anthocyanidin aglycon, decoration with sugar moieties, and transport into the vacuole (**Fig. 2**). Anthocyanidin aglycones are formed from the substrate naringenin-chalcone which is processed by CHI, F3H, DFR, and ANS (Winkel-Shirley, 2001). Alternative steps in the anthocyanidin biosynthesis are catalysed by F3′H and F3’5′H leading to alternative substrate for DFR and ANS, giving rise to structurally different anthocyanidins. Anthocyanidins are converted into anthocyanins through decoration with sugars, catalysed by glycosyltransferases (GTs). GTs can accept a broad range of substrates but modify a specific position of the aglycon (Offen *et al*., 2006; Wang *et al*., 2019; Yi *et al*., 2020). Usually, a 3-O-glycosylation is the first modification step followed by 5-O-glycosylation and possibly additional decoration steps. Anthocyanins are then imported into the vacuole where they are stored. The molecular mechanisms underlying this import remain poorly understood but anthocyanin deficient mutants show that the anthocyanin ‘escort’ protein (ligandin) AN9 (Edwards *et al*., 2000; Mueller *et al*., 2000; Kitamura *et al*., 2004) and MATE and/or ABC transporters are involved in the transport process (Marinova *et al*., 2007; Francisco *et al*., 2013) in model experimental systems.

**Figure 2.**
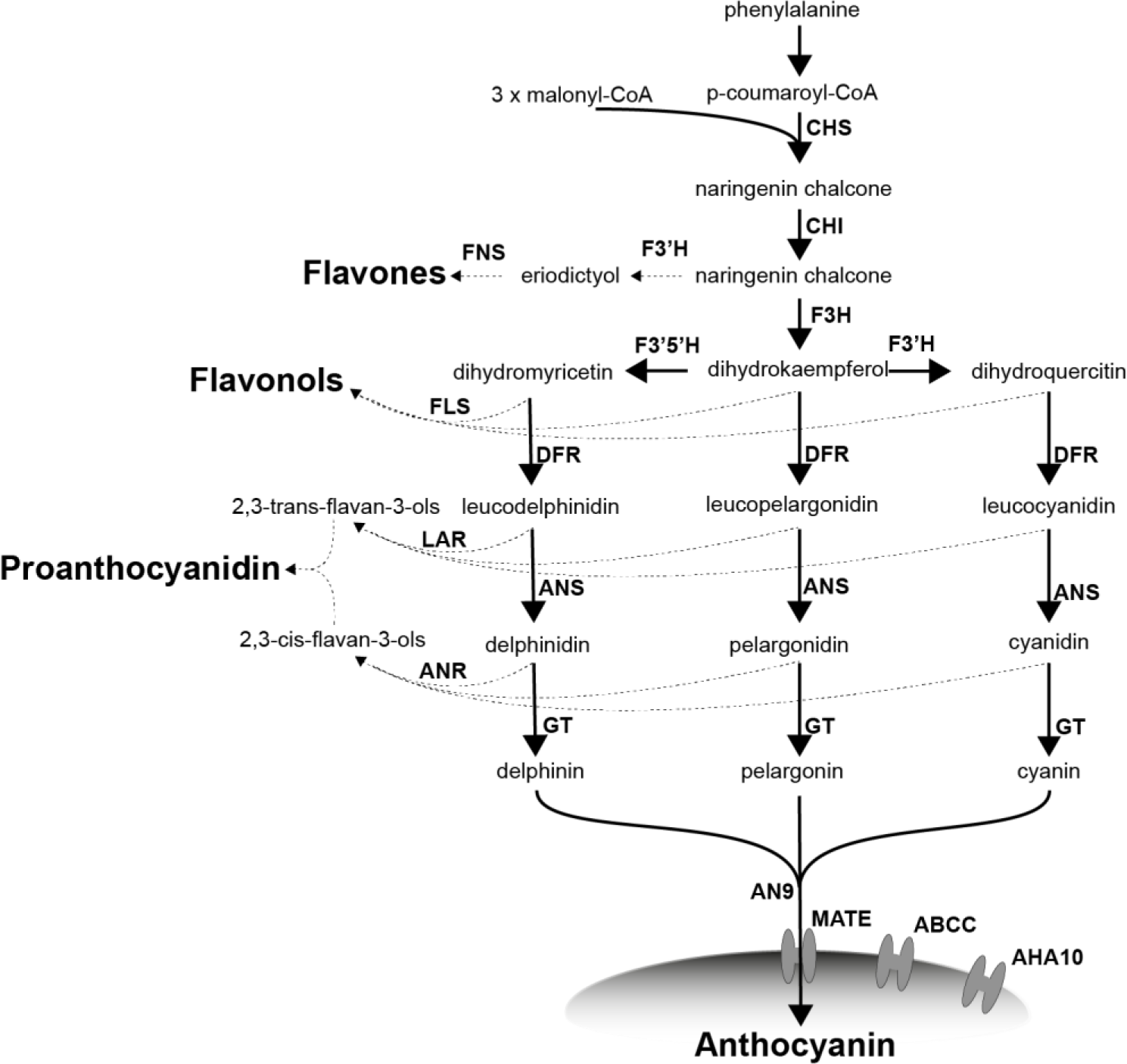
Simplified flavonoid biosynthesis pathway. CHS (naringenin-chalcone synthase), CHI (chalcone isomerase), FNS (flavone synthase), FLS (flavonol synthase), F3H (flavanone 3-hydroxylase), F3′H (flavonoid 3′-hydroxylase), F3′5′H (flavonoid 3′,5′-hydroxylase), DFR (dihydroflavonol 4-reductase), ANS (anthocyanidin synthase), LAR (leucoanthocyanidin reductase), and ANR (anthocyanidin reductase), GT (glycosyltranferase; here the arrow represents glycosyltransferase enzymes in general rather than a specific glycosyltransferase, as glycosylations takes place as series of steps), AN9 (Glutathione S-transferase), MATE (proton antiporter), ABCC (ATP binding cassette protein 1), and AHA10 (Autoinhibited H(+)-ATPase isoform 10). MATE, ABCC, and AHA10 are involved in the anthocyanin transport from the cytoplasm into the vacuole. Shaded oval represents the vacuole in which anthocyanins are stored.

The fate of the anthocyanin synthesis pathway has previously been studied in five betalain-pigmented species in Caryophyllales: *Beta vulgaris* and *Spinacia oleracea* (Amaranthaceae), *Phytolacca americana* (Phytolaccaceae), *Mirabilis jalapa* (Nyctaginaceae), and *Astrophytum myriostigma* (Cactaceae) (Shimada *et al*., 2004, 2005, 2007; Polturak *et al*., 2018; Hatlestad *et al*., 2015; Sakuta *et al*., 2021). Several hypotheses have been explored to explain the lack of anthocyanins in betalain-pigmented Caryophyllales lineages, including: a) loss of anthocyanin synthesis genes, b) loss or changing function of anthocyanin synthesis genes, c) tissue-specific loss of transcriptional activation of anthocyanin biosynthesis gene due to modification to cis-regulatory regions, and d) degeneration of the canonical MBW complex responsible for activation of anthocyanin synthesis genes. To date there is little evidence for wholesale loss of anthocyanin synthesis genes because studies in three separate species have found dihydroflavonol 4-reductase (*DFR*) and anthocyanidin synthase (*ANS*) maintained in three different betalain-pigmented species, *S. oleracea*, *P. americana*, *A. myriostigma*, probably because of their pleiotropic role in proanthocyanidin synthesis. There is conflicting evidence on loss of function of anthocyanin synthesis genes, because canonical gene function for *ANS* and *DFR* is conserved in *S. oleracea* and *P. americana* (Shimada *et al*., 2004, 2005), yet a truncated ANS protein *M. jalapa* lacks anthocyanidin synthase activity suggesting that loss of anthocyanins in *M. jalapa* may be attributable to loss of *ANS* function (Polturak *et al*., 2018). Modification of the cis-regulatory regions of *ANS* and *DFR* has been inferred in some studies but remains inconclusive due to the heterologous nature of promoter binding assays (Shimada *et al*., 2007, Sakuta *et al*., 2021). Finally in both *B. vulgaris* and *A. myriostigma* the trans-acting PAP1 homologs have lost the ability to bind canonical bHLH partners in heterologous assays, which is suggested to contribute to a loss of ability in activating anthocyanin biosynthesis genes *in planta* (Hatlestad *et al*., 2015, Sakuta *et al*., 2021).

The current dominant model for loss of anthocyanin pigmentation assumes the presence of functional anthocyanin synthesis genes, and attributes modification to low gene expression of *DFR* and *ANS* as the key mechanism in anthocyanin loss (Hatlestad *et al*., 2015, Sakuta *et al*., 2021). But this emphasis is influenced by hitherto limited observations on a small number of late-acting components (essentially *DFR* and *ANS*) in the flavonoid synthesis pathway (**Fig. 2**). The exclusive focus on DFR and ANS is problematic, as these enzymes do not catalyse committed anthocyanin biosynthesis steps *per se* and are also involved in the production of proanthocyanidins (**Fig. 2**), which are retained in betalain-pigmented species (Shimada *et al*., 2005). Additionally, few early components of the flavonoid synthesis pathway have been examined except for *CHS*, and absent from consideration are the steps such as glycosylation enzymes and post-synthesis anthocyanin transporters, which are critical for anthocyanin stability and accumulation. Finally, these observations have been made on just five betalain-pigmented species which may not be sufficient to resolve the diversity of mechanisms underlying anthocyanin loss, especially given a hypothesis of multiple transitions to betalain pigmentation.

Here we sought to leverage the recent expansion in genomic and transcriptomic resources to generate a gene-rich and species-rich comparative framework, to revisit the fate of the anthocyanin synthesis pathway in the context of multiple transitions to betalain pigmentation. Specifically, we were motivated by two hypotheses: a) that the mechanisms underlying the loss of anthocyanins may be different across different transitions to betalains, e.g., different genes down-regulated or lost; b) that additional mechanisms are required to explain the potential irreversibility of anthocyanins loss suggested by a scenario of multiple transitions to betalains. Using 3,833 publicly available RNA-seq datasets and genome sequence assemblies, we report on the evolutionary fate and expression profiles of 18 flavonoid pathway genes, across 301 species and 31 families, and representing three of the four putative origins of betalain pigmentation.

## MATERIALS AND METHODS

### Data source and processing raw sequences

Most sequence data used in this study were transcriptome and genome assemblies from the One Thousand Plant Transcriptome (1KP) project and other studies (Matasci et al., 2014; Walker et al., 2018; Pucker et al., 2020a). Additional transcriptome assemblies were generated based on publicly available RNA-Seq datasets of *Halostachys caspica*, *Myosoton aquaticum*, *Oxyria digyna*, *Achyranthes bidentata*, *Dysphania schraderiana*, *Hammada scoparia*, *Hololachna songarica*, and *Gymnocarpos przewalskii* using a previously established protocol (Haak *et al*., 2018). Briefly, this involved trimming with Trimmomatic v0.39 (Bolger *et al*., 2014) followed by an assembly with Trinity v2.4 (Grabherr *et al*., 2011) with k=25 and a prediction of peptide sequences (Haak *et al*., 2018). A total of 361 transcriptome assemblies and 21 genome assemblies of 359 Caryophyllales species were included in the analyses (see data availability statement for details). The completeness of the predicted peptides in transcriptome and genome assemblies was evaluated through the presence of well-conserved Benchmarking Single Copy Orthologs (BUSCOs) with BUSCO v3 (Simão *et al*., 2015), run in protein mode with an e-value cutoff of 1e-3 and considering at most 10 hits on all predicted peptide sets using the ‘embryophyta odb9’ reference gene set (Zdobnov *et al*., 2017).

### Identification of candidate sequences

To perform a comprehensive analysis of the flavonoid biosynthesis, a thorough annotation of sequences in transcriptome and genome assemblies is required. Annotation is based on sequence similarity to previously characterized sequences. Previously characterized protein sequences for each step in the flavonoid biosynthesis (Pucker *et al*., 2020), modification, and transport pathway including CHS, CHI, F3H, F3′H, F3′5 ′H, FLS, DFR, ANS, LAR, ANR, A3GT/UFGT78D2, A5GT/UFGT75C1, F3GT/UFGT79B1, AN9, MATE, AHA10, and ABCC were used as baits (search queries) for the identification of candidate sequences with a high degree of similarity to baits. This collection of bait sequences was further extended by identification of orthologous sequences in datasets representing >120 species of major plant lineages (NCBI and phytozome datasets) based on a previously described approach (Yang *et al*., 2015). Smith-Waterman alignment-based searches with SWIPE v2.0.12 (Rognes, 2011) were conducted against each transcriptome or genome assembly, and up to 100 hits per bait with a minimum bit score of 30 were considered in the initial step and manually refined through iterative construction of gene trees with FastTree2 (Price *et al*., 2010) and removal of sequences on long branches likely to represent distantly-related or non-homologous sequences. Next, the extended set of bait sequences were used to further identify candidate sequences in the Caryophyllales following the same iterative approach (Yang *et al*., 2015). Alignments are inferred with MAFFT v7.475 with default auto settings (Katoh & Standley, 2013). For comparison, the analysis was also performed for the carotenoid biosynthesis pathway, in which *A. thaliana* protein sequences served as baits for the identification of homologs in the Caryophyllales (**Table S1**) based on the phylogenetic approach (Yang *et al*., 2015) as described above. The carotenoid biosynthesis was separately pulled out as a control because it is a pigment pathway yet biochemically distinct and part of the primary metabolism (as opposed to specialised metabolism), so less likely to show a systematic difference (i.e., due to condition-specific lack of expression) between anthocyanin and betalain-pigmented groups.

### Construction of phylogenetic trees

For the construction of gene trees, peptide sequences of outgroup species and Caryophyllales were aligned via MAFFT v7.475 using default auto settings (Katoh & Standley, 2013). Next, the aligned amino acids were substituted with the corresponding codons using pxaa2cdn from phyx (Brown *et al*., 2017). Alignment columns with occupancy below 10% were removed via phyx (Brown *et al*., 2017), (pxclsq −p 0.1). raxml-ng v0.9 (Kozlov *et al*., 2019) was used to generate final trees using the GTR+G model and 100 rounds of bootstrapping. Monophyletic or paraphyletic groups of sequences from a single species’ transcriptome assemblies could represent true paralogs or isoforms and were reduced to one representative sequence using a publicly available script (Yang & Smith, 2014). Briefly, clusters of monophyletic sequences of a single species are identified and reduced to the single longest transcript in the cleaned alignment. Paraphyletic sequences that are at most one node away from the monophyletic cluster were also masked. Trees were visualized in FigTree (http://tree.bio.ed.ac.uk/software/figtree/). Several iterations of tree building, and manual cleaning were performed to generate the final gene trees. For example, exceptionally long branches on isolated sequences can sometimes indicate an alignment or annotation issue which escaped initial filtering, where difficult to explain long branches were recognized, the alignment was manually examined to understand any issues – sequences which were clearly mis-annotated on part of their length or otherwise suspiciously misaligned were manually removed. Additional outgroup sequences were included to distinguish between related gene families: stilbene synthases and other polyketide synthases for CHS, short-chain dehydrogenases for DFR (Moummou *et al*., 2012). Sequences of closely related gene families were investigated in a joined alignment and tree to ensure proper assignment of the candidate sequences. F3’H and F3’5’H were investigated together. F3H, FLS, and ANS were analyzed together to clearly separate these closely related 2-oxoglutarate dependent dioxygenase sequences. We used an overlap-based approach to label duplication nodes in the gene tree, requiring at least two species to overlap between the two daughter clades to map a gene duplication event to a node, and therefore only detect deeper level gene duplication events represented by at least two species in our taxon sampling.

### Quantifying gene expression

We collected a comprehensive set of 4,071 publicly available RNA-Seq datasets of the Caryophyllales (https://github.com/bpucker/CaryoAnthoBlock). While public RNA-Seq datasets are a valuable resource, metadata about the experimental settings can be incomplete or inaccurate e.g., the classification of DNA sequencing data as RNA-Seq. Filtering steps were applied to exclude unreliable datasets. It is well known that a substantial amount of reads in an RNA-seq experiment belongs to a small number of highly abundant transcripts. Assessing this distribution allowed the identification and removal of normalized libraries and other artifacts which would not be suitable for quantitative analyses. The proportion of expression assigned to the 100 most abundant transcripts (top100) was determined for all datasets. Cutoffs were identified based on the distribution of these values. Only datasets with >10% and <80% of the total transcript per million (TPM) assigned to the top100 transcripts were subjected to down-stream analyses. 3,833 RNA-Seq datasets belonging to 301 species passed these filters. Where possible, only paired-end datasets were considered, because these reads can be assigned to similar transcripts with higher confidence. Quantification was performed with sequencing runs as individual data points. Since the number of data sets per species is highly variable, all species are represented by their mean value per gene in downstream analyses to avoid an overrepresentation of species with many available data sets. The available metadata were compared between anthocyanin-pigmented and betalain-pigmented groups to exclude systematic differences (**Table S2**). As each species is represented with a single average value in the comparison between pigmentation groups, the most abundant tissue type was identified for each species. As UTR annotation or representation in a transcriptome assembly is error- prone, only coding sequences were used for the quantification of transcript abundances. kallisto v0.44 was applied with default parameters to quantify read abundance based on paired-end datasets (Bray *et al*., 2016). Since we do not know the fragment size in libraries of single end datasets, an average fragment size of 200bp with a standard deviation of 100bp was assumed for all samples. Individual count tables were merged to generate one table per species and filtered as described above using customized Python scripts (https://github.com/bpucker/CaryoAnthoBlock). Gene expression was compared between anthocyanin-pigmented and betalain-pigmented lineages for all steps in the flavonoid biosynthesis. The sum of the transcript abundances (TPMs) of all isoforms of a gene were added up per RNA-seq sample (**Fig. S4**). Isoforms are all sequences that were phylogenetically assigned to the same function in the pathway through the steps described above. The combination of large numbers of datasets generated for different tissues under various conditions results in a high level of noise. However, only strong biological signal should emerge from the broad-scale comparative analysis, yet precise quantifications among different lineages are not feasible. The average value representing each species comprises a species-specific number of samples that have different degrees of diversity, therefore, we refrained from displaying the variation of this data sets in a single value.

### Micro-synteny

To clarify if the absence of *AN9* is due to a lack of transcription in the studied samples or due to gene loss in the betalain-pigmented species, the genome sequences of four representative Caryophyllales species were analyzed. Since the physical location of *AN9* is known in *Solanum lycopersicum*, it was possible to identify the syntenic region in the genome sequences of *Vitis vinifera* and Caryophyllales species. *Beta vulgaris* (betalain transition 2, B2 for short here after), *Dianthus caryophyllus* (anthocyanin-pigmented), *Mesembryanthemum crystallinum* (B3), and *Carnegiea gigantea* (B4) represent different lineages of the core Caryophyllales (see **Fig. 1**). Unfortunately, no genome sequence is available for one betalain lineage (Stegnospermataceae, B1). Collinear regions that lack *AN9* but harbour the flanking genes were inspected to search for *AN9* to examine if there was any evidence or pseudogenisation in process or if the genes had been lost in their entirety. Microsynteny around the *S. lycopersicum AN9* locus was analysed via JCVI using mcscan and the synteny function (Tang *et al*., 2008). The -cscore cutoff was set to 0.1 to ensure high sensitivity and only the most likely region was considered. This approach relies on a BLAST-based comparison of genes in the compared species but chains adjacent BLAST hits to detect collinear blocks of genes between two genomes. Consequently, this syntenic analysis is more reliable than a simple search based on sequence similarity alone, by focusing the search on the likely region of a gene’s location and detecting similar sequences which are syntenically conserved and thus more likely to be truly homologous.

## RESULTS

### Most flavonoid pathway genes were detected across all betalain-pigmented families except for *F3’5’H* and *AN9*

We searched for the following 18 genes in the flavonoid pathway within our transcriptome and genome sequence assemblies, and performed phylogenetic analyses to explore relationships, duplication, and loss: chalcone synthase (*CHS*), chalcone isomerase (*CHI*), flavanone 3-hydroxylase (*F3H*), flavonoid 3’-hydroxylase (*F3*’*H*), flavonoid 3’,5’-hydroxylase (*F3*’*5*’*H*), flavonol synthase (*FLS*), flavone synthase (FNS), dihydroflavonol 4-reductase (*DFR*), anthocyanidin synthase (*ANS*), leucoanthocyanidin reductase (*LAR*), anthocyanidin reductase (*ANR*), anthocyanidin 3-O-gucosyltransferase (*A3GT*), anthocyanidin 5-O-gucosyltransferase (*A5GT*), flavonoid 3-O-gucosyltransferase (*F3GT*), glutathione S-transferase 26 (*AN9/TT19*), proton antiporter (*MATE/TT12*), ATP binding cassette protein 1 (*ABC*), and autoinhibited H(+)-ATPase isoform 10 (*AHA10/TT13*). The bulk of datasets used in this analysis are transcriptomic in origin, and can only offer proof of gene presence, as apparent gene absence may simply be due to lack of expression. However, coupled with annotated genome assemblies representing three of the inferred origins of betalain pigmentation (*Beta vulgaris*, *Mesembryanthemum crystallinum*, and *Carnegeia gigantea*), the combined genomic and transcriptomic datasets are informative with respect to the broad scale patterns of low gene expression and/or loss (**Fig. 3)**.

**Figure 3.**
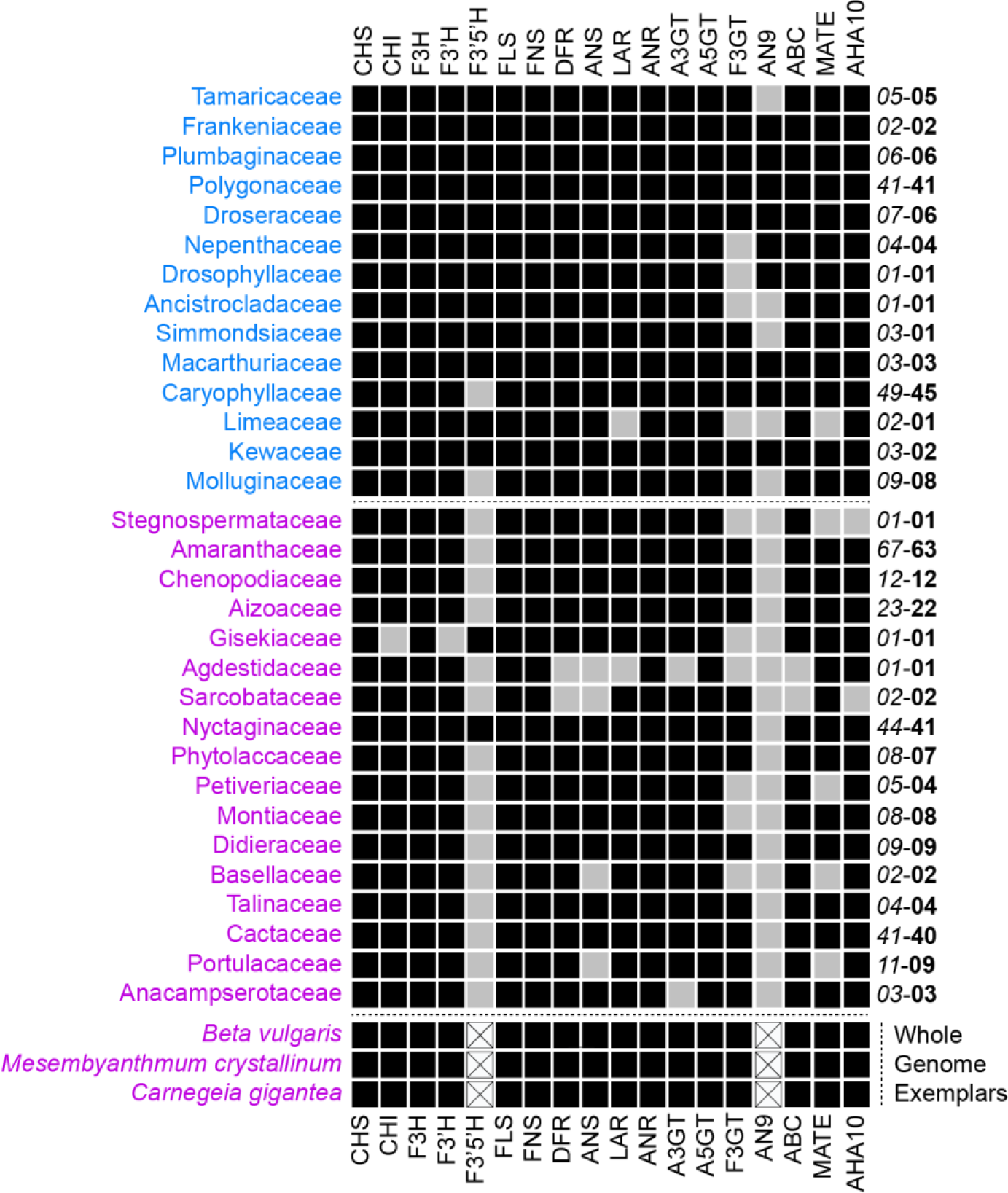
Detection of flavonoid biosynthesis genes in 359 Caryophyllales species summarized at the family level. Families are sorted by pigmentation state into anthocyanin-and betalain-pigmented (blue=anthocyanin, pink=betalain) to highlight the consistent differences between pigment types. Generally, most genes of the flavonoid biosynthesis are present in most families. Only *F3′5 ′H* and *AN9* are consistently missing from betalain-producing families. Species with exceptionally well annotated contiguous genome sequences that represent the three betalain origins were included at the bottom in italics to add additional support to the pattern. *CHS* (naringenin-chalcone synthase), *CHI* (chalcone isomerase), *FNS* (flavone synthase), *FLS* (flavonol synthase), *F3H* (flavanone 3-hydroxylase), *F3′H* (flavonoid 3′-hydroxylase), *F3′5′H* (flavonoid 3′,5′-hydroxylase), *DFR* (dihydroflavonol 4-reductase), *ANS* (anthocyanidin synthase), *LAR* (leucoanthocyanidin reductase), and *ANR* (anthocyanidin reductase), *GT* (glycosyltranferase), *AN9* (glutathione S-transferase), *MATE* (proton antiporter), *ABC* (ATP binding cassette protein 1), and *AHA10* (autoinhibited H(+)-ATPase isoform 10). Black=presence in at least one transcriptome or genome assembly in the family, grey=not detected in transcriptome assembly, white with a cross=absence unable to detect in whole genome sequencing data. Number on the right-hand side indicate number of transcriptome and genome assemblies sampled (*italics)* and number of species (**bold**).

We focused on evidence for deeper level gene loss with the gene data summarized at the level of family, in line with data on pigment status. In some anthocyanin-pigmented families we detected occasional sporadic gene absence without apparent phylogenetic pattern for: *F3H, F3′5 ′H, LAR, F35GT, AN9* and *MATE*. These apparent gene absences in anthocyanic taxa usually appeared in lineages with very little transcriptomic coverage and therefore higher probability of stochastic lack of detection. In general, most flavonoid biosynthesis, decoration, and transport associated genes are maintained and expressed in betalain-pigmented families But in some betalain-producing families that lack whole genome data, we were unable to find transcriptomic evidence for the following genes: *F3GT*, *MATE*, and *AHA10* in Stegnospermataceae, *CHI*, *F3H*, and *F3′H* in Gisekiaceae; *CHI*, *DFR*, *ANS*, *LAR*, and *ANR* in Agdestidaceae; *CHI*, *F3H*, *DFR* and *ANS* in Sarcobataceae; *ANS* and *LAR* in Basellaceae; *ANS*, *LAR* and *ANR* in Portulacaceae. However, Stegnospermataceae, Gisekiaceae, Agdestidaceae and Sarcobataceae are all monotypic families, comprising only a single species, and represented by a single transcriptome assembly in our analyses, again representing a higher probability of lack of detection **(Fig. 3)**.

Putative stochastic absences aside, two stronger patterns of gene absence emerge in relation to betalain-pigmentation lineages. First, we find no evidence for the presence of F3′5 ′H in 15 out of 17 betalain-pigmented families including in genome assemblies from *B. vulgaris*, *M. crystallinum*, and *C. gigantea*. We recovered a striking pattern of repeated absence from transcriptome and genome assemblies for the anthocyanin carrier protein AN9 in betalain-pigmented lineages that could be explained by gene loss (**Fig. 3**). 5 tree of *AN9* (**Fig. 4a**), we recovered numerous sequences from anthocyanic non-core Caryophyllales species and core Caryophyllales anthocyanin-pigmented Caryophyllaceae, Macarthuriaceae and Kewaceae. Importantly, *only* anthocyanin-pigmented species are represented in this tree, and no sequences were detected from a betalain-pigmented species. A further screen of the highly contiguous genome sequences of betalain-pigmented species did not reveal any *AN9* sequences. To rule out any mis-annotation issues, based on the *Solanum lycopersicum AN9* ortholog (Solyc02g081340), we identified the corresponding micro-syntenic regions in the genome sequences of betalain-pigmented species. Although the region shows conserved microsynteny in the flanking sequences, we did not find a sequence or fragments of a sequence with significant similarity to *AN9* in betalain-pigmented *Beta vulgaris*, *Mesembryanthemum crystallinum*, or *Carnegiea gigantea* (**Fig. 4b**). These species represent independent betalain-pigmented lineages, and our phylogenetic reconstruction support that *AN9* has been separately and completely lost in multiple betalain lineages (**Fig. 4c**). The probability of missing *AN9* by chance in all betalain-pigmented families (0/17), assuming that the proportion of absence in anthocyanin-pigmented families (5/14) represents the probability of failing to detect *AN9* when it is present, would be below 0.001 (binomial probability). This estimation does not account for the better representation of betalain-pigmented species datasets within families.

**Figure 4.**
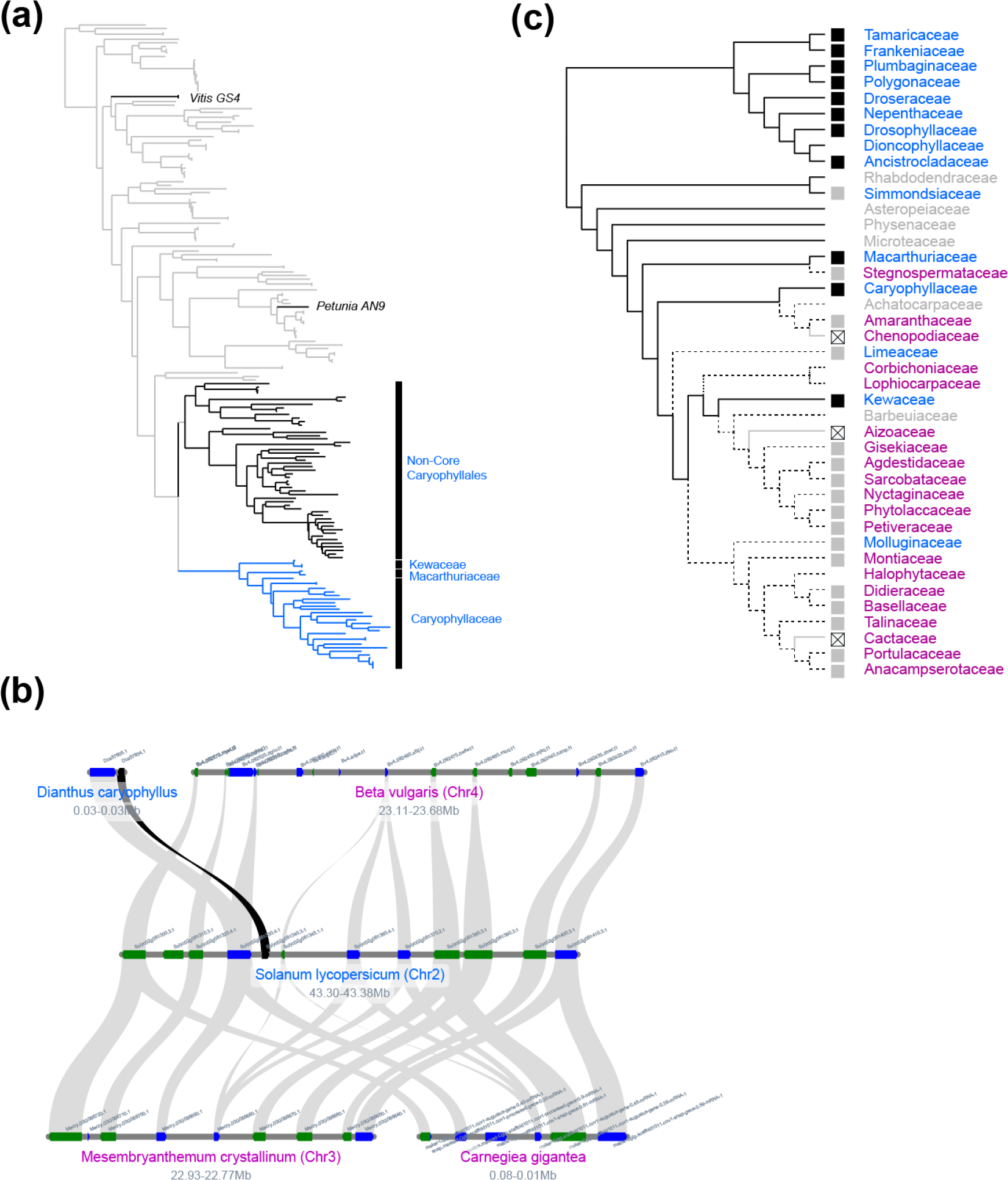
Loss of *AN9* homologs in betalain-pigmented lineages. (a) A phylogenetic analysis revealed the presence of *AN9* homologs in most anthocyanin-pigmented Caryophyllales species, but the absence from all betalain-pigmented species in 31 families sampled. (grey = non-Caryophyllales outgroups, black = non-core anthocyanic Caryophyllales, blue = anthocyanic core Caryophylales). Functionally characterised outgroup orthologs *Vitis GS4* and *Petunia AN9*, which are known to have anthocyanin transport activity, are labelled on the tree. (b) Microsynteny analysis of the *AN9* locus (black) of genome sequences representing an anthocyanin-pigmented outgroup (*Solanum lycopersicum*) and anthocyanin-pigmented in-group (*Dianthus caryophyllus)* and three betalain-pigmented species (*Beta vulgaris*, *Mesembryanthemum crystalinum*, *Carnegiea gigantea*) supports gene loss in the betalain-pigmented lineages (dark blue=gene on forward strand, green=gene on reverse strand, black line indicates position and synteny of AN9 homolog between *Solanum lycopersicum* and *Dianthus caryophyllus*). (c) Parsimony-based reconstruction of *AN9* loss assuming losses are irreversible, and with the conservative assumption that absence of a gene from the transcriptome is not proof of absence. Black lines = presence, grey lines = absence, dotted lines = ambiguous, blue=anthocyanin, pink=betalain, gray box = no detected, black box = gene detected crossed box = not detected in genome, no box = missing data.

### Flavonoid biosynthesis gene trees show extensive gene duplication across core Caryophyllales, including in betalain-pigmented lineages

Based on the phylogenetic topologies for each of the 18 flavonoid synthesis genes (**Fig. S1**), we observed that the flavonoid pathway within the Caryophyllales is shaped by patterns of repeated gene duplications (**Fig. 5**). Notably, many duplications occur within betalain-pigmented lineages and are maintained over relatively long periods of evolutionary time. Overall, *CHS* shows one of the most dynamic patterns with a duplication event early in core Caryophyllales, prior to the divergence of *Macarthuria*, and numerous family specific duplications within the anthocyanic Caryophyllaceae, betalain-pigmented Amaranthaceae s.l. and Cactaceae, and multiple rounds of duplications within the betalain-pigmented Nyctaginaceae (**Fig. S1**). *FNS* is widely duplicated in multiple betalain lineages including Nyctaginaceae. *F3*′*H* is duplicated in the anthocyanic Caryophyllaceae, the betalain-pigmented Didieraceae, and has undergone two rounds of duplication within the betalain-pigmented Nyctaginaceae. *DFR* duplicated in Nyctaginaceae and Polygonaceae. *ANS* has duplicated in Polygonaceae, Nyctaginaceae and the Portulacineae alliance. As is evident from the above description, almost the entire flavonoid biosynthesis pathway is maintained and went through gene duplication in the betalain-pigmented Nyctaginaceae. *CHS*, *CHI*, *F3H*, *F3*′*H*, and *ANS* were duplicated within Nyctaginaceae, corresponding to a whole genome duplication event at the base of the tribe Nyctagineae (Yang *et al*., 2018); and *DFR* shows a duplication in the common ancestor of *Mirabilis* and *Commicarpus* in Nyctaginaceae. Given the patterns of gene family evolution, we note that full length *ANS* genes are in fact maintained across the Nyctaginaceae, including in *Mirabilis jalapa*. The identification of paralogous copies of *ANS* in Nyctaginaceae more broadly, and *Mirabilis jalapa* specifically, is significant because an earlier study (Polturak *et al*., 2018) suggested that truncation and loss of function of one of the *ANS* copies in *Mirabilis jalapa* may underlie loss of anthocyanin pigmentation in this species. Our findings indicate however that *Mirabilis jalapa* retains a full length *ANS* sequence, in addition to the truncated copy (**Fig. S2**).

**Figure 5.**
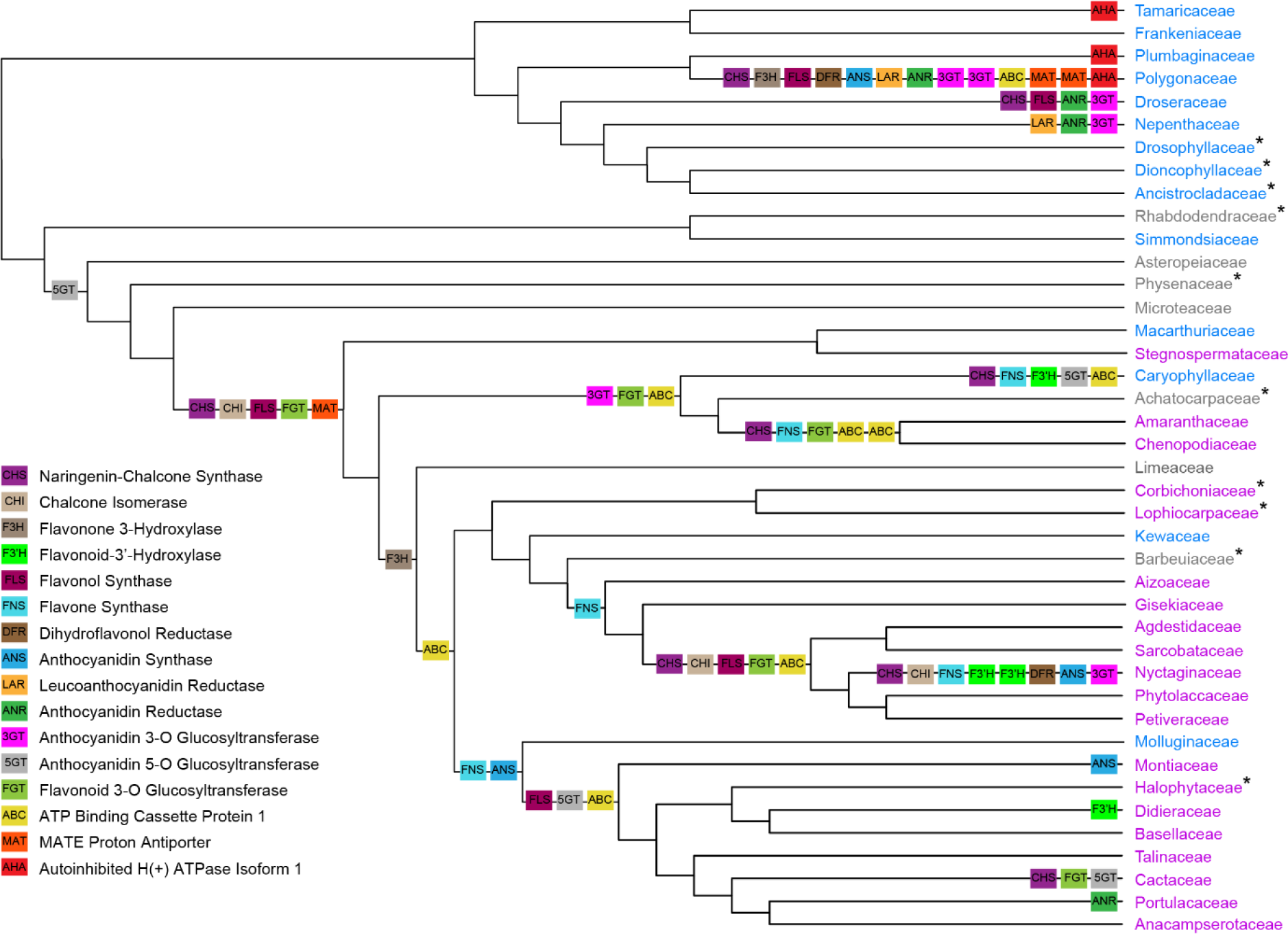
Summary of flavonoid biosynthesis gene duplications in the Caryophyllales. Gene duplication events for flavonoid biosynthesis genes are mapped to a family-level phylogeny based on Walker *et al*., 2018 and Sheehan *et al*., 2020. The representation is restricted to deeper level gene duplications and excluded events below the genus level. CHS (naringenin-chalcone synthase), CHI (chalcone isomerase), FNS (flavone synthase), FLS (flavonol synthase), F3H (flavanone 3-hydroxylase), F3′H (flavonoid 3′-hydroxylase), F3′5′H (flavonoid 3′5′-hydroxylase), DFR (dihydroflavonol 4-reductase), ANS (anthocyanidin synthase), LAR (leucoanthocyanidin reductase), and ANR (anthocyanidin reductase), 3GT (anthocyanidin 3-O-gucosyltranferase), 5GT (anthocyanidin 5-O-glucosyltranferase), FGT (flavonoid 3-O-glycosyltransferase), AN9 (glutathione S-transferase), MAT (proton antiporter), ABC (ATP binding cassette protein 1), and AHA (autoinhibited H(+)-ATPase isoform 10). Family names in blue = anthocyanin, pink = betalain, grey = unknown pigmentation status. Asterisks indicate families which are not well represented in the analyzed data set.

### Many late-stage flavonoid biosynthesis genes show reduced transcript abundance in betalain-pigmented species compared to anthocyanin-pigmented species

The recent accumulation of transcriptomic datasets enabled the systematic comparative investigation of transcript abundances for all flavonoid biosynthesis genes, and the broad comparison of transcript abundances between anthocyanic and betalain-pigmented species (**Fig. 6**). Here, through a large-scale data mining of 4,071 publicly available RNA-seq datasets representing 301 species across Caryophyllales we observed a generally reduced transcript abundance in most genes in the flavonoid biosynthesis pathway in betalain-pigmented versus anthocyanin-pigmented species. This observation was very common, but the differences are far more dramatic for some genes than others.

**Figure 6.**
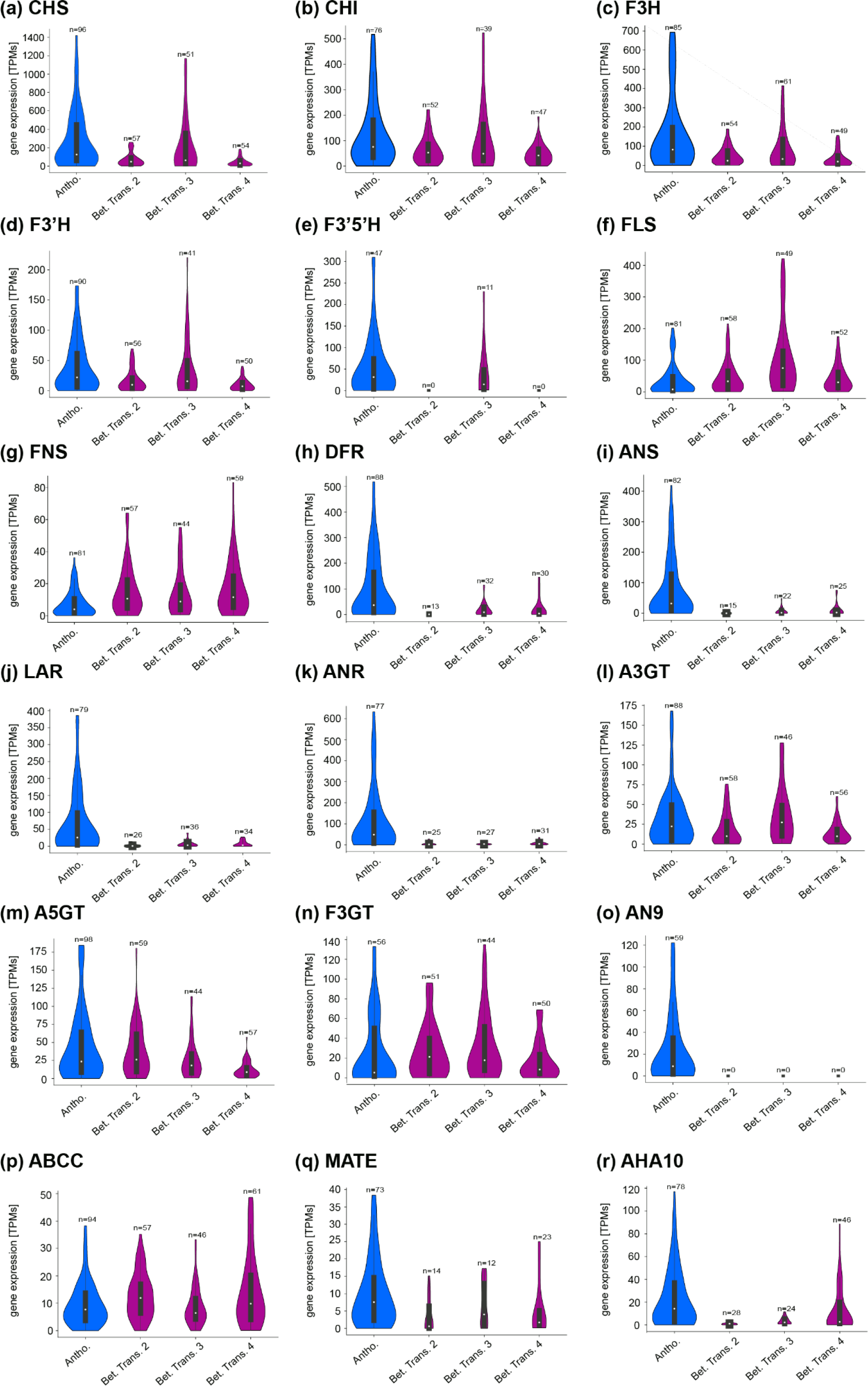
Comparative analysis of flavonoid biosynthesis gene expression in the Caryophyllales. The gene expression in anthocyanin-pigmented species (blue) is compared to the gene expression in species of three betalain transitions (magenta) (see **Fig 1b** for illustration of three origins). (a) CHS (naringenin-chalcone synthase), (b) CHI (chalcone isomerase), (c) F3H (flavanone 3-hydroxylase), (d) F3′H (flavonoid 3′-hydroxylase), (e) F3′5′H (flavonoid 3′5′-hydroxylase), (f) FLS (flavonol synthase), (g) FNS (flavone synthase), (h) DFR (dihydroflavonol 4-reductase), (i) ANS (anthocyanidin synthase), (j) LAR (leucoanthocyanidin reductase), (k) ANR (anthocyanidin reductase), (l) A3GT (anthocyanidin 3-O-gucosyltranferase), (m) A5GT (anthocyanidin 5-O-glucosyltranferase), (n) F3GT (flavonoid 3-O-glycosyltransferase), (o) AN9 (glutathione S-transferase), (p) MATE (proton antiporter), (q) ABCC (ATP binding cassette protein 1), and (r) AHA10 (autoinhibited H(+)-ATPase isoform 10). Blue=anthocyanin-pigmented lineages, pink=betalain-pigmented lineages.

This pattern is apparent for some early acting components (*CHS*, *CHI*, *F3H*, *F3′H*, & *F3′5 ′H*) but is especially pronounced for the late acting components (*DFR*, *ANS*, *LAR*, *ANR*, *MATE*, & *AHA10*). This phenomenon was clearly visible in data representing the three putative betalain origins that were sampled. An analysis investigating the transcript abundance of carotenoid biosynthesis genes did not reveal similar differences between anthocyanin-pigmented species and species of the three putative betalain origins (**Fig. S3**), suggesting that the pattern is not due to the heterogeneity of publicly available RNA-seq data we used. Although expression of later-acting flavonoid biosynthesis genes leading to anthocyanins and proanthocyanidins are highly reduced in betalain species, several genes acting in other branches of the flavonoid biosynthesis show little reduction. For example, the flavonol biosynthesis gene *FLS* shows almost no difference between anthocyanic and betalain-pigmented lineages. Although *CHS* transcript was observed at substantially lower abundance in betalain species, its abundance is still relatively high compared to other genes in the pathway, implying a substantial production of the key flavonoid substrate naringenin chalcone in betalain-pigmented lineages.

## DISCUSSION

Despite the loss of anthocyanins, the presence of a broad range of other flavonoids (flavones, flavonols and proanthocyanins) is well documented in betalain-pigmented species (Iwashina, 2015). The branched nature of the flavonoid synthesis pathway (Ho & Smith, 2016; Ng *et al*., 2018) means that most enzymatic steps attributed to anthocyanin synthesis are pleiotropic with respect to these other flavonoids (Fig. 1a). Consequently, most studies have found that late acting enzymes in the anthocyanin synthesis pathway are functionally maintained in betalain-pigmented lineages (Shimada *et al*., 2004, 2005, 2007; Sakuta *et al*., 2021). Given the proposed maintenance of these enzymes, loss of anthocyanins has instead been mostly attributed to regulatory changes in the expression of late acting enzymes. Here we have sought to test this model across the entire flavonoid pathway, using phylogenetically dense genomic scale datasets, and across multiple origins of betalain pigmentation. As explored in the remainder of the discussion, we find that important aspects of this model hold true, with two developments; a) we find evidence of wholesale and repeated loss of two significant genes within the anthocyanin synthesis pathway within betalain pigmented lineages, and; b) we find some evidence for reduced transcription, not just of late-acting anthocyanin synthesis genes, but across the majority of genes within the flavonoid biosynthetic pathway.

### Duplication and loss of flavonoid biosynthesis genes in Caryophyllales

Of the 18 components of the flavonoid pathway examined here, almost all are broadly conserved and actively expressed in betalain-pigmented families (**Fig. 3**), consistent with the reported presence of flavonols, flavones, and proanthocyanidins in both betalain - pigmented and anthocyanin-pigmented species (Iwashina, 2015). In addition, we found extensive evidence of gene duplication of flavonoid pathway genes, across core Caryophyllales, and notably also within betalain-pigmented lineages (**Fig. 5**). Many of these paralogous genes persist over considerable evolutionary time, suggesting their maintenance via sub- or neo-functionalisation. Gene duplication is a well described phenomenon with respect to the flavonoid pathway (Yang *et al*., 2002; Yonekura-Sakakibara *et al*., 2019; Piatkowski *et al*., 2020) but the level of gene duplication in Caryophyllales suggests an evolutionary dynamism in flavonoid biosynthesis to a degree that is perhaps unanticipated in betalain-pigmented lineages. Different genes showed different numbers of duplication events, with the early acting *CHS* showing the highest degree of duplication whereas fewer gene duplication events were detected in the late-acting *DFR* and *ANS*. Extensive duplication across multiple flavonoid genes is also occurring in a lineage-specific fashion, in some cases clearly associated with whole genome duplication events, as are documented for Nyctaginaceae (Yang *et al*., 2015, 2018). On the one hand, the loss of anthocyanins and an apparent shift to tyrosine-dominant metabolism (see below; Lopez-Nieves *et al*., 2018) suggests that we should not anticipate functional radiation in flavonoid metabolism. On the other hand, many Caryophyllales are found in highly abiotically stressful environments, and the further evolution of non-anthocyanin flavonoids may have occurred in response to this. Additionally, some duplications are likely maintained merely due to neutral fixation or dosage effects.

Recently truncation and loss of ANS activity in the flavonoid biosynthesis gene *ANS* has been invoked as a potential mechanism to explain loss of anthocyanins in the betalain-pigmented species *M. jalapa* (Polturak *et al*., 2018). All genes of the anthocyanin synthesis pathway are expressed in the flowers of *M. jalapa*, and yet no anthocyanins are produced. This was attributed to a deletion in a florally expressed *MjANS* (Polturak *et al*., 2018) as *MjANS* is unable to complement an *Arabidopsis thaliana ans* mutant (Polturak *et al*., 2018). However, we find evidence of an *ANS* gene duplication, which has given rise to two clades within Nyctaginaceae, both containing full length *ANS* variants but with one clade also containing the truncated version previously detected in *M. jalapa*. (**Fig. S2**). Both full length and truncated variants are present in *Mirabilis jalapa*. Based on these data, we suggest that wholesale or functional loss of *ANS* is unlikely to underlie the loss of anthocyanins in *Mirabilis*. In this study we find examples of other lineages in which *ANS* may be absent, but these examples are based on limited transcriptomic data from monotypic lineages, with the interesting exception of the Portulacaceae, which is well represented by transcriptome assemblies. The lack of detection of *ANS* across multiple transcriptome samples within Portulacaceae may merit further investigation, but in general, we show that most betalain-pigmented species retain a full length *ANS* gene, and we find no genomic evidence for *ANS* gene loss or loss of ANS function (at least by clear frame-shifting or long indels) in annotated genome sequences of betalain-pigmented species.

F3′5 ′H and F3′H are enzymes acting at branch points within the flavonoid biosynthesis pathway, catalyzing the conversion of dihydrokaempferol to dihydromyricetin or dihydroquercitin, respectively. *F3′H* and *F3′5′H* are both Cytochrome P450 enzymes and form two sister subfamilies CYP75A and CYP75B, respectively (Yonekura-Sakakibara *et al*., 2019). Both subfamilies are deeply conserved across flowering plants, with *F3′5′H* recruited from *F3′H* before the divergence of angiosperms and gymnosperms. However, we were unable to detect the presence of *F3′5 ′H* CYP75A lineage in the transcriptomes of 17/23 families within core Caryophyllales, including 15/17 betalain-pigmented families. Furthermore, we were unable to detect the *F3′5 ′H* CYP75A in all three annotated genome sequences from betalain-pigmented species (**Fig. 3**). Dihydromyricetin, the product of *F3′5 ′H* activity, can be converted either to myricetin-derived flavonols, or alternatively, is the key substrate in the pathway leading to the blue anthocyanin delphinin. *F3′5′H* has previously been documented to be rapidly pseudogenised and deleted in anthocyanin-pigmented species that have transitioned away from delphinin-based blue towards red coloured flowers (Smith & Rausher, 2011; Wessinger & Rausher, 2014), indicating a major role for *F3′5′H* in the production of blue anthocyanins (Ho & Smith, 2016). Interestingly, in the extensive documentation of flavonoids across Caryophyllales (Iwashina, 2015), quercetin-type flavonoids derived via *F3′H* enzymatic activity are very common, but myricetin-type flavonoids derived via F3′5′H enzymatic activity are correspondingly extremely rare, supporting the general absence of *F3′5′H* activity in Caryophyllales. It is unclear to what extent the loss of *F3′5′H* is related to the evolution of betalain pigments, but blue flowers are rare across Caryophyllales, including in the anthocyanin-pigmented Caryophyllaceae, perhaps resulting in the widespread loss of *F3′5′H*. However, the presence of F3′5′H in two nested betalain lineages does imply that the absence of F3′5′H in certain lineages might be explained by repeated reduction of expression in the studied tissues or loss that has occurred repeatedly and towards the tips of the phylogeny rather than as a single early-occurring evolutionary event.

The AN9 family of glutathione S-transferases (which includes the *AN9* gene in *Petunia hybrida* and the *TT19* ortholog in *A. thaliana*) are thought to be an important component in anthocyanin transport and accumulation (Mueller *et al*., 2000). Mutants of *AN9*/*TT19* are deficient in anthocyanin accumulation, and evidence from *A. thaliana*, indicates that TT19 acts as a transport-associated protein (van Houwelingen *et al*., 1998; Kitamura *et al*., 2004). Anthocyanin accumulation without TT19 was only observed in plants with a substantially increased metabolic flux in the flavonoid biosynthesis (Jiang *et al*., 2020), which is the opposite of our observations in the Caryophyllales. The current model proposes that TT19 binds and stabilizes anthocyanins, and potentially shuttles them from the cytoplasm to the tonoplast, where they are acylated and transported into the vacuole (Sun *et al*., 2012). Given the importance of AN9 for anthocyanin accumulation, it is striking that *AN9* orthologs are completely absent from all transcriptome assemblies and annotated genome sequences in betalain-pigmented species yet are detectable in three anthocyanin lineages within core Caryophyllales. The fact that *AN9* is detected in Kewaceae, Caryophyllaceae and Macarthuriaceae, indicates it was retained from their common ancestor as a plesiomorphic state. On the assumption that lost *AN9* loci cannot be regained, we inferred multiple losses of *AN9* orthologs, and suggest that *AN9* has been lost independently in at least three of our putative betalain origins (**Fig 4c)**. Apparent sporadic lack of detection of *AN9* from anthocyanin-pigmented families can best be explained by the small number of available transcriptome assemblies for these lineages.

We are unable to determine with the current data whether the loss of *AN9* homologs is responsible for the initial loss of anthocyanins, especially given alternative mechanisms such as reduced expression of *ANS* and *DFR*, and the potential deprivation of related transcription factors (Hatlestad *et al*., 2015; Sakuta *et al*., 2021). Nonetheless the loss of *AN9* is significant for our understanding of directionality in pigment evolution. Previously, the maintenance but restricted expression of flavonoid synthesis genes, *ANS* and *DFR*, in the proanthocyanidin-containing seed coats of betalain-pigmented species gives a clear evolutionary mechanism for multiple reversals back to anthocyanin pigmentation from a betalain ancestor (**Fig. 1a)**, i.e., restoring expression patterns of *ANS* and *DFR* in the shoot could restore anthocyanin biosynthesis, assuming presence of all other components of the pathway. However, a recent study has shown that restoration of anthocyanin pigmentation in betalain-pigmented *A. myriostigma* is possible by genetic engineering. Heterologous expression of *DFR* and *ANS*, and separately, heterologous expression of *Arabidopsis PAP1* (the canonical trans-activator of *DFR* and *ANS)* in *A. myriostigma* requires heterologous expression of *PhAN9* to cause anthocyanin pigmentation (Sakuta *et al*., 2021). Although Sakuta *et al*. did not identify that the native *AN9* had been lost in betalain lineages, clearly *AN9* is implicated as decisive factor for potential anthocyanin synthesis in betalain-pigmented species, not solely the expression of *DFR* and *ANS*. Crucially, the repeated losses of *AN9* from betalain-pigmented lineages, recovered in this study, imply repeated anthocyanin loss in core Caryophyllales, consistent with the previous finding of repeated specialisation to betalain pigmentation (Sheehan *et al*., 2020).

### Reduced expression of multiple flavonoid biosynthesis genes in betalain-pigmented Caryophyllales

In advance of any discussion of our comparative expression analyses, we acknowledge their limitations. On the one hand, like many bioinformatic reanalyses, the data we interrogated were not originally acquired with our goals and analyses in mind. But on the other hand, these publicly available transcriptomes represent a remarkably broad species and tissue sampling that is beyond the scope of a single study. Nonetheless, there are disparities in the number of transcriptome datasets available among species, and lack of consistency between species in terms of sampling across different tissue types, developmental stages, and stress treatments. In absence of any corresponding metabolite data, we are unable to correlate gene expression patterns with flavonoid metabolites of interest. Furthermore, because of repeated gene duplication events, and in absence of functional data for different paralogs, many of which are newly identified in this study, we were forced to integrate expression values across multiple paralogs. We are re-assured that the macroevolutionary patterns we report are not the consequence of systematic bias, because single genes of the flavonoid biosynthesis like *FLS* and the analysis of the analogous carotenoid pigmentation pathway reveal no systematic differences in expression. As broad-brush strokes, these analyses provide an important but largely qualitative insight into flavonoid pathway gene expression, which must be interpreted with caution.

In general, we observe lower expression of most anthocyanin pathway genes in betalain versus anthocyanin-pigmented species, across the three inferred betalain origins studied here. This pattern is apparent for some early acting components (*CHS*, *CHI*, *F3H*, *F3′H*, & *F3′5 ′H*) but is especially pronounced for the late acting components (*DFR*, *DFR*, *LAR*, *ANR*, *MATE*, & *AHA10*). This low transcript abundance is consistent with previous studies that found that loss of anthocyanin pigmentation is associated with cis- and/or trans-regulatory changes to enzymatic genes (Shimada *et al*., 2004, 2005, 2007; Sakuta *et al*., 2021). Several flavonoid biosynthesis genes do not fit this pattern of low transcriptional expression in betalain-pigmented lineages including *FLS* and *FNS*, and the three genes encoding glycosylation enzymes, here termed *A3GT*, *A5GT* and *F3GT*. Along with the control analysis of the carotenoid pathway, this indicates that the patterns of reduction we do observe are not the result of some artefactual and systematic bias in the datasets. The glycosylation enzymes have been previously described as being broadly promiscuous (Offen *et al*., 2006; Wang *et al*., 2019; Yi *et al*., 2020) and some have been shown to have the ability to decorate betalains (Vogt *et al*., 1999; Vogt, 2002). Given this substrate promiscuity, it is perhaps not surprising that we observe little difference in the expression levels of these enzymes in anthocyanin-pigmented versus betalain-pigmented species. Finally, we can attribute comparable expression levels of *FLS* and *FNS* in betalain-pigmented versus anthocyanin-pigmented species, to the continued presence of flavonols and flavonones in betalain-pigmented species. Perhaps the redirection of the bulk of flavonoid substrates to flavonols and flavonones, instead of anthocyanins, is reflected in the continued high expression *FLS* and *FNS*.

The overall trend of reduction in expression across the pathway, including early-acting genes, is interesting, given that betalain-pigmented species do continue to produce flavonols and flavonones. One explanation for the apparent overall reduction in the expression of flavonoid biosynthesis genes is the notion of a shift from phenylalanine-derived metabolism to tyrosine-derived metabolism within core Caryophyllales (Lopez-Nieves *et al*., 2018). A gene duplication in the arogenate dehydrogenase lineage, has given rise to a novel isoform of arogenate dehydrogenase (*ADH◻*), which has lost feedback sensitivity, and which increases tyrosine production at the expense of phenylanine production in heterologous assays in *N. benthamiana* (Lopez-Nieves *et al*., 2018). The evolution of the *ADH◻* isoform therefore could potentially limit the availability of phenylalanine, which might therefore be reflected in the lower gene expression levels in flavonoid pathways. Alternatively, simply the absence of anthocyanin as an end-product, might mean there is less demand for naringenin chalcone entering flavonoid metabolism, which is again reflected in generally lower gene expression levels. This is consistent with previous studies that have found that early-acting genes in the anthocyanin pathway and their regulators are targets for selection when there are evolutionary transitions in total amount of anthocyanin production (Jung *et al*., 2009; Payyavula *et al*., 2013; Tian *et al*., 2017).

## Conclusion

Given a working hypothesis of multiple shifts to betalain pigmentation, we re-visited mechanisms for anthocyanin loss. With respect to our original hypotheses, we find little evidence that the mechanisms of anthocyanin loss are different between different betalain origins. Across all three betalain origins we see a similar and marked low transcript abundance of many flavonoid genes, and especially a similar severe loss of expression of the more committed genes for anthocyanin synthesis, *DFR* and *ANS*, and the apparent wholesale loss of AN9. But given the crude nature of our analyses, we cannot discriminate the order of change, whether loss of *DFR* and *ANS* expression preceded or followed loss of *AN9*. It is also unclear whether it is cis- or trans-regulatory change, or a combination, that underlies the reduced expression of these genes, therefore remains possible that with closer interrogation the genetic mechanisms underlying loss of *DFR* and *ANS* expression in different origins may be distinct. Given that recent evidence shows ectopic expression of *AN9* is critical for the genetic engineering of anthocyanin biosynthesis in betalain-pigmented lineages (Sakuta *et al*., 2021), it is intriguing that this loss of *AN9* has apparently happened convergently in multiple betalain-pigmented lineages, consistent with the hypothesis of multiple origins of betalain pigmentation.

## ACKNOWLEDGEMENTS

We thank members of the Brockington lab for discussion and careful reading of the manuscript. We thank the Center for Biotechnology (CeBiTec) at Bielefeld University and de.NBI for providing an environment for computational analyses. We thank Benoit van der Rest for sharing a collection of SDR sequences and Andrea Berardi for helpful discussion. We thank all colleagues and the wider community who performed sequencing studies in the Caryophyllales and shared raw data that enabled this analysis. We acknowledge support from the following funding bodies: BP, Deutsche Forschungsgemeinschaft (DFG, German Research Foundation) – 436841671; NWH, Woolf Fisher Cambridge Scholarship; JCC, DOE, GSP DE-SC0008834; SFB, BBSRC High Value Chemicals from Plants Network; AC, SFB &YY, National Science Foundation NSFDEB-NERC award #1939226.

## AUTHOR CONTRIBUTION

The work was conceived by BP and SFB. Unpublished genomic resources were provided by WCY and JC. Analyses were conducted by BP with support of NWH, AC, and YY. Figures were prepared by SFB and BP. The manuscript was written by SFB and BP. All authors read and approved the manuscript.

## DATA AVAILABILITY

RNA-seq data sets analyzed in this study are available at the SRA/ENA. A list of the analyzed data sets, FASTA files containing bait sequences and sequences identified in this study, and Python scripts developed for this study are available at github: https://github.com/bpucker/CaryoAnthoBlock.

## SUPPLEMENTARY INFORMATION

**Fig. S1** Phylogenetic trees of the flavonoid biosynthesis and flavonoid transport genes.

**Fig. S2** Analyses of *Mirabilis jalapa ANS* gene copies.

**Fig. S3** Carotenoid biosynthesis gene expression analysis

**Fig. S4** Illustration of the cross-species gene expression calculation that forms the basis of Fig. 6.

**Table S1** Peptide sequences of carotenoid biosynthesis genes that were used to identify homologs in the Caryophyllales.

**Table S2** Comparison of RNA-seq tissue types between anthocyanin-pigmented and betalain-pigmented plants.

**Table S3** Analysis of the genomic region where AN9 would be expetected in betalain-pigmented species.

