## Supplementary material for "Evolutionary blocks to anthocyanin accumulation and the loss of an anthocyanin carrier protein in betalain-pigmented Caryophyllales": Fig. S1

The following Supporting Information is available for this article:

**Fig. S1** Phylogenetic trees of the flavonoid biosynthesis and flavonoid transport genes.

**Fig. S2** Analyses of *Mirabilis jalapa ANS* gene copies.

**Fig. S3** Carotenoid biosynthesis gene expression analysis

**Fig. S4** Illustration of the cross-species gene expression calculation that forms the basis of Fig. 6.

**Table S1** Peptide sequences of carotenoid biosynthesis genes that were used to identify homologs in the Caryophyllales.

**Table S2** Comparison of RNA-seq tissue types between anthocyanin-pigmented and betalain-pigmented plants.

**Table S3** Analysis of the genomic region where AN9 would be expetected in betalain-pigmented species.

**Fig. S1** Phylogenetic trees of genes in the flavonoid biosynthesis and flavonoid transport steps were constructed by aligning peptide sequence with MAFFT, replacing amino acids with the corresponding codons, trimming to a minimal alignment column occupancy of 10%, inference of phylogeny with RAxML, and rooting by non-Caryophyllales outgroups. Blue = anthocyanin-pigmented lineages, pink = betalain-pigmented lineages

**[See PDF]**

**Fig. S2** Comparison of truncated and full length ANS sequences in *Mirabilis jalapa*. (a) The phylogenetic positions of ANS(1) and ANS(2) indicates a gene duplication in the Nyctaginaceae. ANS sequences are highlighted in red. Two ANS clades are distinguished by color: blue and pink, respectively. (b) The full length and the truncated ANS differ by a large gap in an import ANS domain as previously described by Polturak et al., 2018.

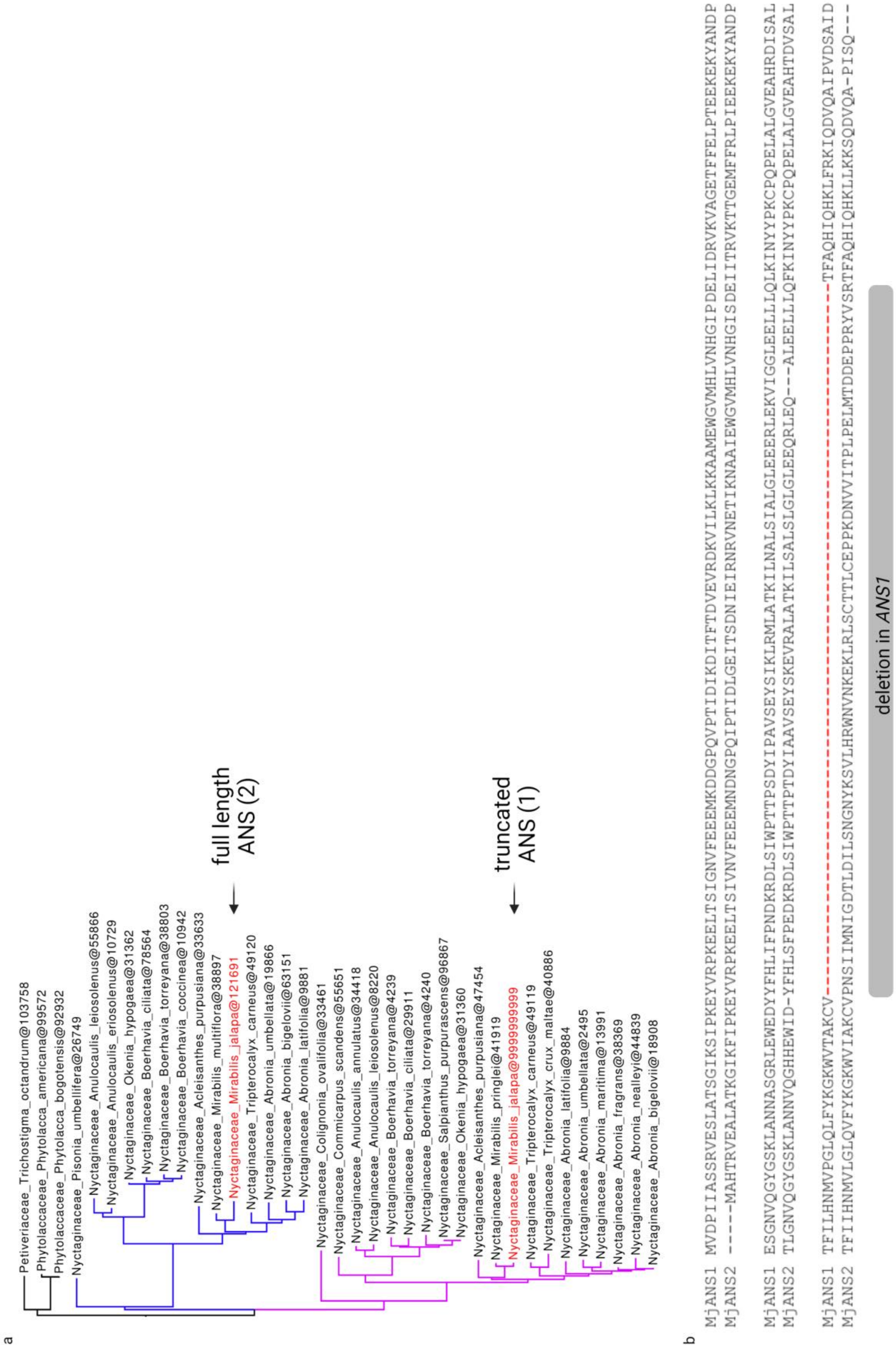

**Fig. S3** Analysis of the transcript abundances of carotenoid biosynthesis genes in anthocyanin-pigmented lineages and the different betalain origins. Please see Table S1 for details about the genes. The position of different transcript abundance plots approximates the position of the encoded enzymes in the pathway.

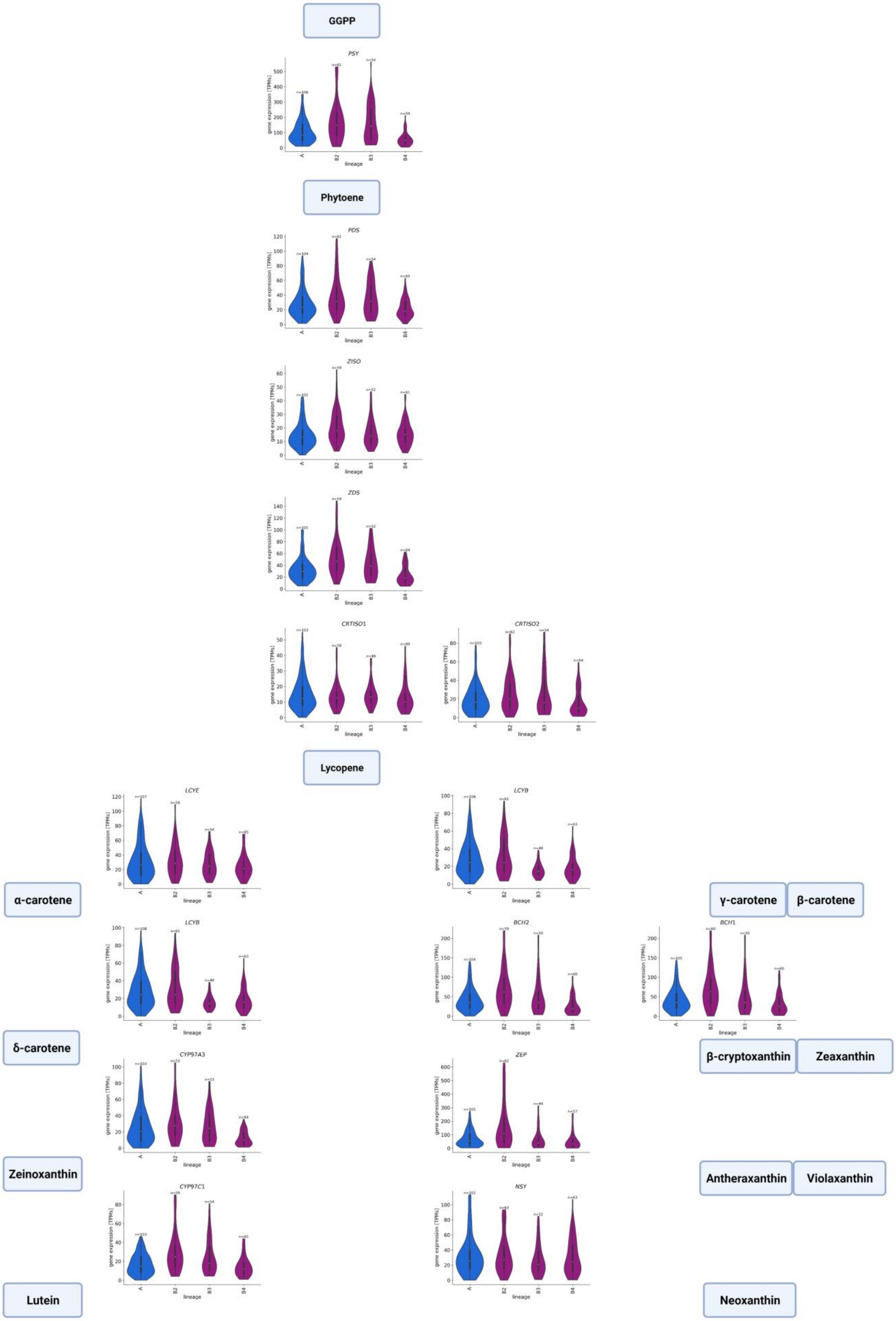
GGPP
PSY
Phytoene
PDS
ZISO
ZDS
CRTISO1
CRTISO2
Lycopene
LCYE
LCYB
α-carotene
γ-carotene
β-carotene
LCYB
BCH2
BCH1
δ-carotene
β-cryptoxanthin
Zeaxanthin
CYP97A3
ZEP
Zeinoxanthin
Antheraxanthin
Violaxanthin
CYP97C1
NSY
Lutein
Neoxanthin
gene expression [TPMs]
lineage
A
B2
B3
B4

**Fig. S4** Illustration of the cross-species gene expression calculation that forms the basis of Fig. 6. This illustrations explains how the values of one gene (geneX) are calculated. All isoforms of one gene are identified across all the analyzed species. For each sample, the abundances of all isoforms of a gene are added up (4+4=8, for sample 1 of species1). The resulting values of all samples are used to calculate an average value per species (8+4+8+6=26, 26/4 = 6.5, for species 1). The values of all anthocyanin-pigmented species are displayed in the blue violin plot and the values of all betalain-pigmented species are displayed in the magenta violin plot.

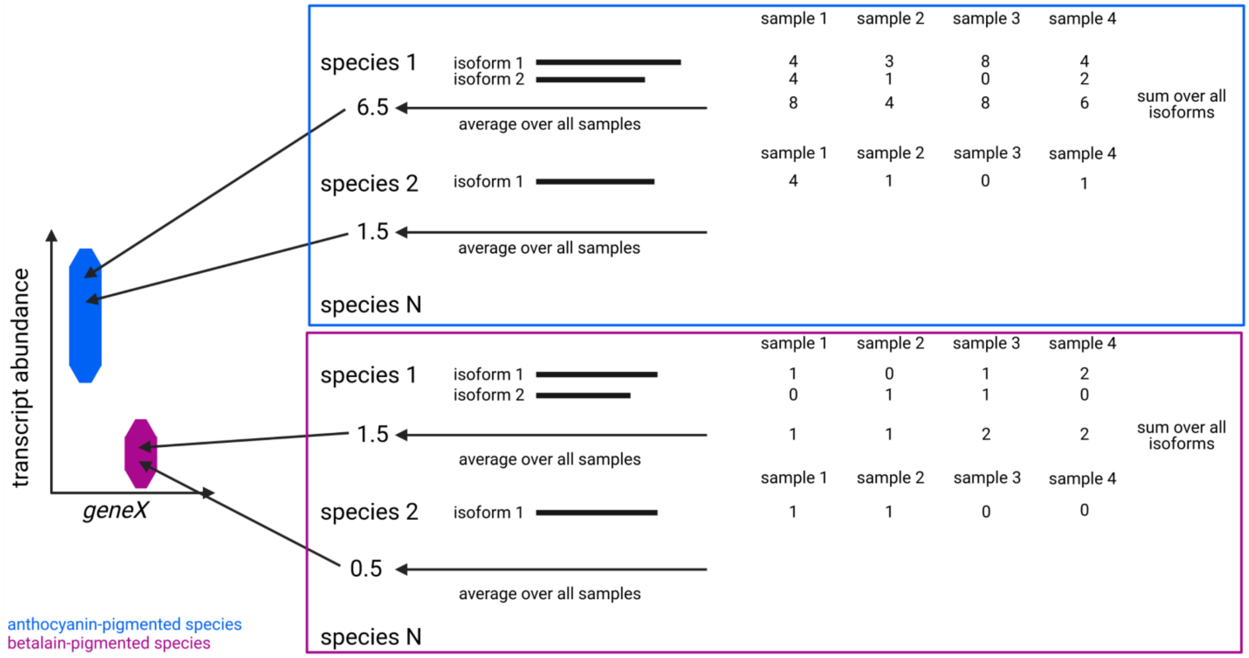


**Table S1** *Peptide sequences of carotenoid biosynthesis genes that were used to identify homologs in the Caryophyllales.*

| **Gene name** | **Arabidopsis Gene Identifier of peptide sequences** |
|---|---|
| PSY | AT5G17230 |
| PDS | AT4G14210 |
| ZDS | AT3G04870 |
| ZISO | AT1G10830 |
| CRTISO1 | AT1G06820 |

| | |
|---|---|
| CRTISO2 | AT1G57770 |
| LCYB | AT3G10230 |
| LCYE | AT5G57030 |
| BCH1 | AT5G25700 |
| BCH2 | AT5G52570 |
| ZEP | AT5G67030 |
| NSY | AT1G67080 |
| VDE | AT1G08550 |
| CYP97A3 | AT1G31800 |
| CYP97C1 | AT3G53130 |
| CYP97B3 | AT4G15110 |

**Table S2** Comparison of RNA-seq tissue types between anthocyanin-pigmented (anthocyanin) and betalain-pigmented (betalain) plants. The different groups used for the lineage specific comparison are all anthocyanin-pigmented species (A), betalain origin 2 (B2), betalain origin 3 (B3), and betalain origin 4 (B4). The metadata available for the analyzed samples were screened for frequently studied tissue types (leaf/leaves, flower, stem, root, seedling, seed).

| **Group** | **Leaf** | **Flower** | **Root** | **Seedling** | **Stem** | **Seed** | **unknown** |
|---|---|---|---|---|---|---|---|
| anthocyanin | 34 | 8 | 2 | 0 | 0 | 2 | 63 |
| betalain | 73 | 14 | 3 | 6 | 4 | 3 | 83 |
| A | 34 | 8 | 2 | 0 | 0 | 2 | 63 |
| B2 | 25 | 1 | 1 | 4 | 1 | 3 | 25 |
| B3 | 30 | 7 | 1 | 0 | 0 | 0 | 19 |
| B4 | 18 | 6 | 1 | 2 | 3 | 0 | 39 |

**Table S3** *Analysis of the genomic region where AN9 would be expected in betalain-pigmented species. The number of gaps, their total size and the size of the region enclosed by the two flanking genes with a syntenic match in Vitis vinifera are listed.*

| **Betalain origin** | **Species** | **Number of gaps** | **Total gap size** | **Size of region** |
|---|---|---|---|---|
| B2 | *Beta vulgaris* | 11 | 26407 bp | 163936 bp |
| B2 | *Amaranthus hypochondriacus* | 0 | 0 | 23372 bp |
| B2 | *Chenopodium palustre* | 0 | 0 | 115421 bp |
| B3 | *Mesembryanthemum crystallinum* | 0 | 0 | 33122 bp |
| B4 | *Carnegia gigantean* | 3 | 306 | 25490 bp |
| B4 | *Hylocereus undatus* | 1 | 10 | 18109 bp |
| B4 | *Portulaca amilis* | 6 | 510 | 83328 bp |
